# Plumage color evolves as distinct dorsal and ventral modules in Australasian honeyeaters

**DOI:** 10.1101/2022.08.23.504757

**Authors:** Nicholas R. Friedman, Vladimír Remeš

## Abstract

Many animals exhibit contrast between their dorsal and ventral coloration. If selection acts differently on dorsal versus ventral coloration, these body parts should evolve as independent modules of trait evolution, irrespective of ancestral covariance. Here, we compare the evolution of feather color across 11 body regions for a clade of Australasian songbirds (Meliphagoidea). We find evidence for three modules of covarying color regions: dorsal, ventral, and flight feathers. Among these modules, ventral feathers had color that was highly labile, evolving at 3 times the rate for dorsal plumage and 20 times the rate for flight feathers. While both dorsal and ventral plumage tend to be correlated with precipitation or the degree of vegetation, we find that dorsal plumage is twice as similar to colors of background substrates in satellite photos. This finding, which a direct effect of climate in Gloger’s rule does not predict, adds support for background matching as an explanation for geographic gradients in animal color. Furthermore, it suggests that selection for background matching has had a greater effect on dorsal plumage than ventral plumage color. Overall, differential selection on ventral and dorsal colors likely maintains these as distinct modules over evolutionary time scales – a novel explanation for dorsoventral contrast in pigmentation.

## Introduction

Many animals exhibit different coloration on their dorsal side than their ventral side. While several mechanisms for this pattern are shared ancestrally among bilaterians and are critical for development (1), uniformly pigmented species have evolved in many taxonomic groups. Thus, the evolutionary maintenance of this predominant color pattern requires explanation. The proximate developmental mechanisms for contrasting dorsal and ventral pigmentation are well known (2, 3). However, ultimate explanations of the function and phylogenetic history of dorsoventral color contrast remain lacking (4). This study aims to clarify to what extent ventral and dorsal pigmentation tend to evolve under the same or different evolutionary processes.

In birds, plumage coloration plays a central role in mate choice and territoriality, as well as crypsis. Each of these functions has been studied exhaustively in isolation (5). But together, where there is an inherent conflict between advertisement and concealment of presence, these functions have received less attention (6). Dorsoventral color contrast is often explained through optimization of a single function, as in countershading (7). However, we propose that this pattern could also arise as a consequence of different selection pressures (8) for advertisement and concealment acting on dorsal versus ventral coloration.

If coloration on all regions of the body is affected similarly by selection for advertisement and concealment, we should expect them to evolve as a single integrated module (9), with similar rates and associations with the environment. In contrast, if dorsal and ventral coloration are influenced by different evolutionary processes, we should expect them to evolve as distinct evolutionary modules, with different rates and associations with the environment (10). To test these predictions, we compared 8879 reflectance spectra from 11 feather patches across 198 species in a clade of Australasian songbirds, the honeyeaters and allies (Meliphagoidea).

## Results & Discussion

Plumage patches on the dorsal side of specimens we measured tended to be twice as dark as patches on the ventral side (lightness or achromatic brightness; phylogenetically corrected paired t-test, p < 0.05; Figure 1A). Likewise, color contrast among patches (measured as Euclidean distance in PC space, see 11) was on average 2.27 times greater for ventral plumage than for dorsal plumage (p < 0.005). These differences are consistent with differential selection on dorsal and ventral plumage color. However, color differences alone are not sufficient to infer the evolutionary processes that maintain them.

**Figure 1.**
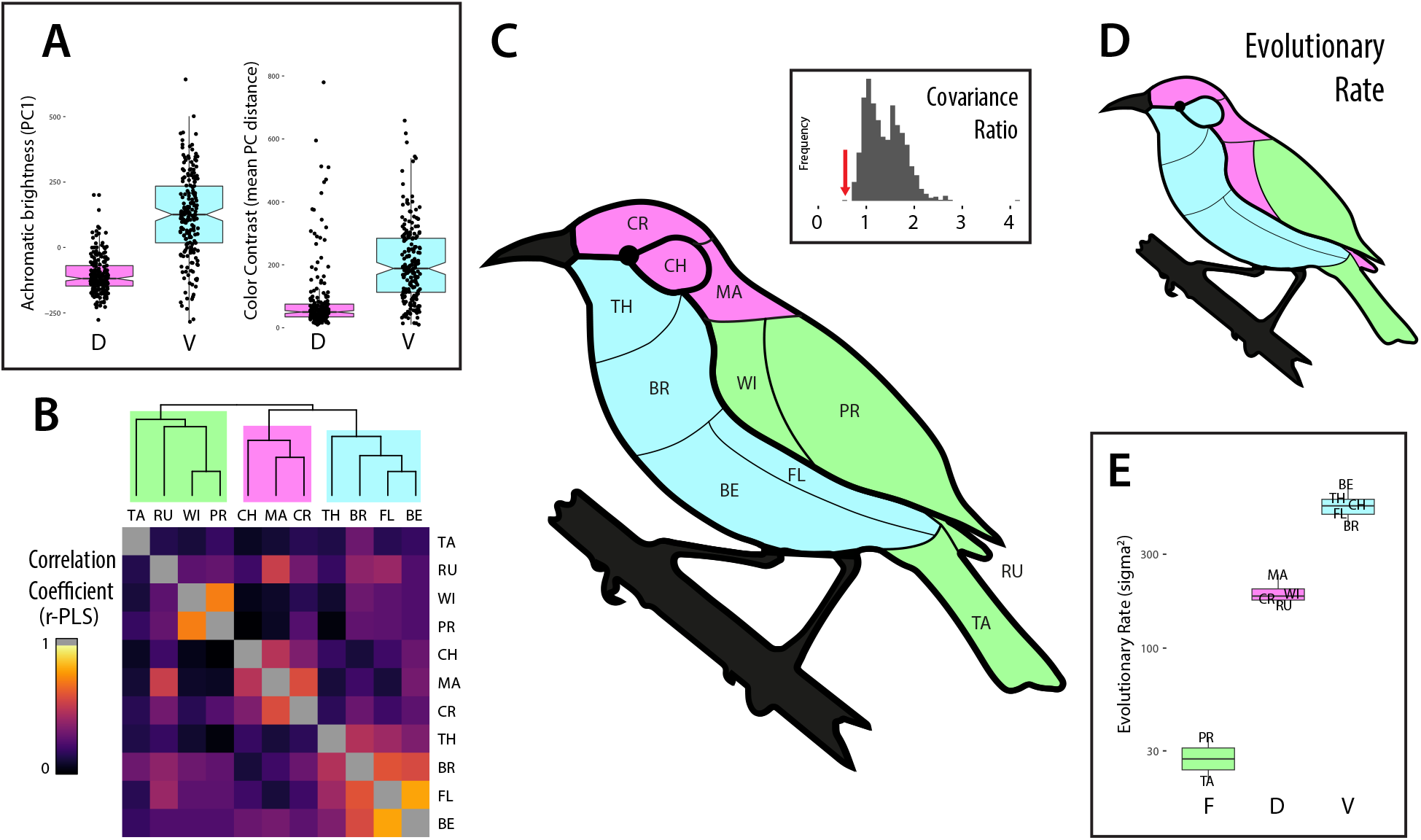
(A) Comparison of dorsal and ventral values for lightness (left) and chromatic contrast among patches, showing that ventral patches are on average brighter and more internally contrasting with other plumage than dorsal patches. (B) Matrix of correlation coefficients for phylogenetic partial least squares comparisons between patches, showing three distinct modules supported by hierarchical clustering. (C) Morphogram of groupings identified in previous panel, showing ventral, dorsal and flight feather modules. Inset describes the observed covariance ratio (red arrow) compared to a simulated null distribution depicted in grey. (D) Morphogram of groupings identified by estimation of joint evolutionary rates (E) supported by kmeans clustering. n = 198 species, 186 for B, C.

To investigate the evolutionary processes underlying plumage color, we estimated the phylogenetically corrected covariance between patches to determine their extent of evolutionary integration. We found that covariance coefficients tended to unite patches into three groups: ventral contour feathers, dorsal contour feathers, and flight feathers with coverts (Figure 1B). Plumage colors on the same or adjacent feather tracts tended to evolve in concert, except when one was on the ventral side and the other dorsal. A covariance ratio test (12) using this module configuration showed greater within-module vs. among-module integration than expected by chance (CR = 0.56, p < 0.001 from 1000 permutations; Figure 1C). Together, these observations indicate the presence of modules exhibiting high evolutionary integration, and suggest that different evolutionary processes influence the independent trajectories of dorsal and ventral plumage.

Elaborate feather color is often used in displays aimed at potential mates or competitors and tends to evolve faster in males than females (5, 13). This implies that rapidly evolving color traits may be influenced by sexual selection pressures that are greater or more variable in their direction. To compare evolutionary rates across plumage patches, we estimated a joint rate across principal components of patch color (14). These rate estimates showed rapid evolution of ventral plumage compared to dorsal or flight feathers. Clustering indicated three modules for dorsal, ventral, and flight feathers, a similar pattern to evolutionary covariance above (Figure 1D). The ratio of evolutionary rates between dorsal and ventral modules was greater for achromatic than chromatic components of color, but the rate order among modules remained consistent (Table S1; Ventral > Dorsal >> Flight). This pattern shows that dorsal and ventral plumage evolve at different rates, and is consistent with (though it does not demonstrate) a greater role for sexually selected signaling functions in ventral plumage than in dorsal plumage (15, 16).

To test the extent to which neutral evolutionary processes can explain these patterns, we conducted simulations with Brownian motion models to produce characters under different rates. We found that high evolutionary rates are likely to produce internally contrasting plumage (Figure S1A) as well as dorsal/ventral brightness contrast (Figure S1B), however such contrast is equally likely in both directions (i.e., ventral should be darker half the time.

We sought to address the difference in evolutionary rates and covariance between ventral and dorsal plumage by investigating the ultimate mechanisms of dorsal plumage evolution. Previous studies have identified a repeated tendency for animals to exhibit darker pigmentation in warmer and more humid regions – an ecogeographic pattern known as Gloger’s Rule (17, 18). Using the same study system, we have shown previously that desert birds tend to have lighter plumage, and that this effect is greater for dorsal than ventral plumage (19). Here as well, we found that birds inhabiting regions with more vegetation (estimated using the Normalized Difference Vegetation Index; NDVI) have darker plumage, especially on their dorsal side (phylogenetic generalized least squares: p < 0.001, R2dorsal = 0.16, R2ventral = 0.10; 20, 21).

While direct climatic explanations for Gloger’s Rule make no predictions about dorsal versus ventral plumage (17), an indirect explanation mediated by selection for background matching does predict this pattern. Thus, we tested whether dorsal plumage color is more similar to background substrates than ventral plumage color. We compared color distances between each patch to pixels in satellite imagery of the specimen’s locality (Figure 2A). Using a phylogenetically corrected generalized linear mixed model (22), we found that dorsal and flight feathers have half the color distance against the background as compared to ventral plumage color (p < 0.001; Figure 2B). This shows evidence for background matching as a functional explanation for variation in dorsal pigmentation, and indicates that indirect rather than direct effects of climate explain the relationship between dark plumage and wet, vegetated areas. Another contributing explanation for the geographical pattern observed may be that in arid climates selection favors individuals better capable of reducing heat gain from solar radiation on their dorsal side (23–25).

**Figure 2.**
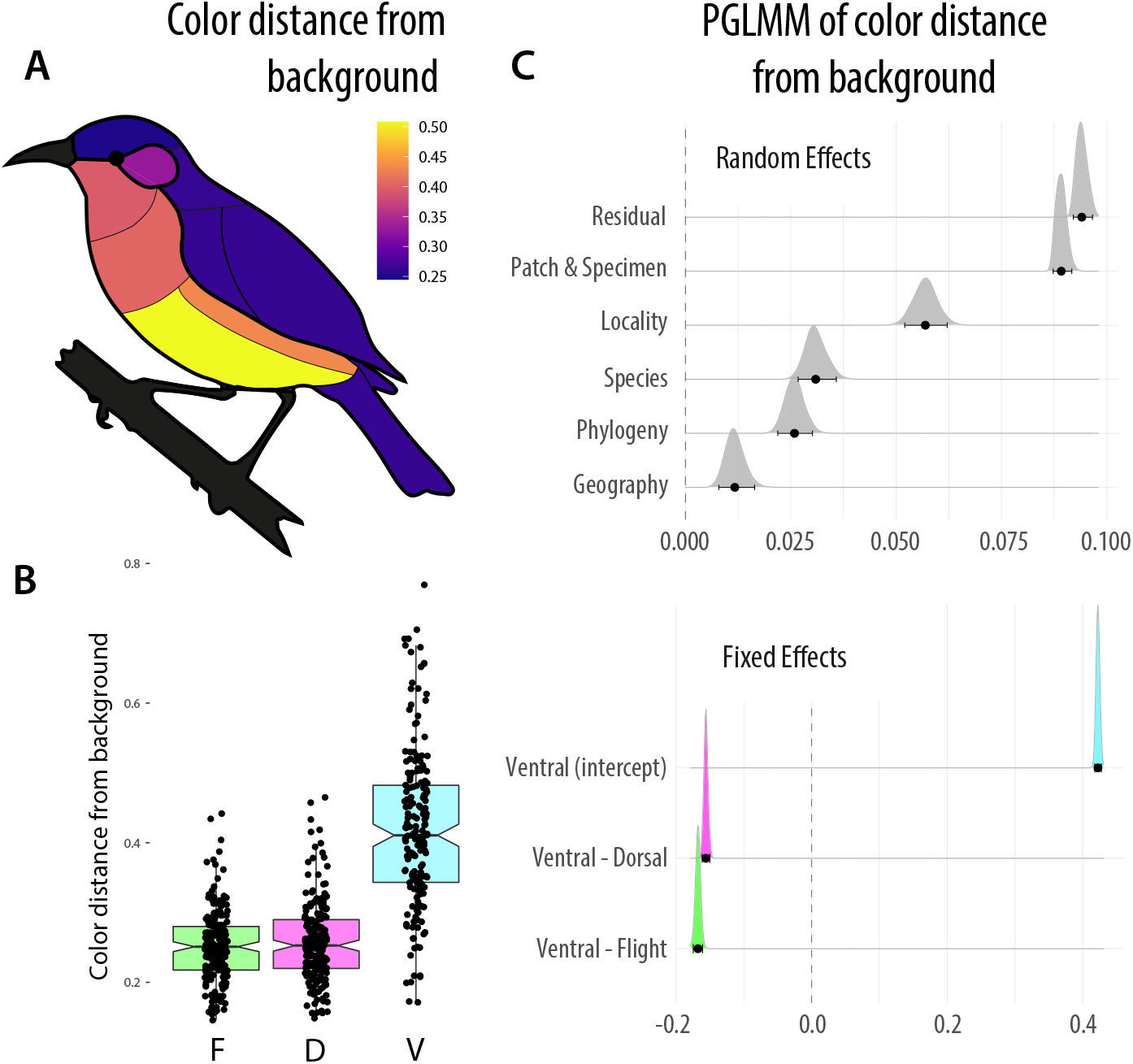
Distance between plumage color and satellite-measured habitat color compared across body regions (A) and previously defined modules (B). Ventral plumage color is more different from the background than the other two modules, also when correcting for phylogenetic and geographic non-independence in a phylogenetic generalized mixed model (C). Effects sizes for random effects are shown above and for fixed effects below, the latter as the ventral intercept and difference compared to it. Mean color distances are shown, n = 196 species.

That colors should differ in function depending on their location is intuitive: the dorsal side typically faces visual predators above while the ventral side faces potential competitors or mates at a similar or lower height (26–28). Likewise, conserved melanism of flight feathers has been reported previously and is likely an adaptation to reduce feather wear (29, 30). Several recent studies have improved our understanding of the evolution of color on different parts of the body (27, 28). However, to our knowledge, no previous studies have reported that dorsal, ventral, and flight feather color can evolve as different modules, at different rates, and under the influence of different processes. We do so here for a clade of songbirds with diverse habitats, visual systems, and mechanisms of coloration (31). Consequently, selection for countershading as a single function (32) may not be necessary to explain the evolution of dorsoventral color contrast; differences in the degree of selection and constraint acting on dorsal and ventral modules can also explain this widespread pattern in animal pigmentation.

## Materials and Methods

We measured plumage colors for 198 species in the honeyeaters and allies using reflectance spectrometry (AvaSpec-2048, Ava-Light-XE, WS-2; Avantes B.V., Apeldoom, Netherlands), and these measurements have been published previously (11, 13, 19). We performed these measurements using vouchered specimens of 3-5 males in the Australian National Wildlife Collection (Specimen Appendix), and averaged three replicate spectra for each of 11 distinct feather patches (illustrated in Figure 1C). We reduced the dimensionality of color variation using a principal components analysis conducted using the covariance matrix of reflectance values in 1nm bins, which is appropriate for phylogenetic comparative methods (33). As is typically the case, PC1 described lightness. We estimated the degree of color contrast for each pair of plumage patches by taking their average pairwise Euclidean distance in PC space (first 10 PCs, including >99% of variance explained) as in a previous study (13) and averaged these for dorsal and ventral patches separately. To analyze achromatic and chromatic color separately, we used PC1 and PCs 2-10, respectively (Figure S2).

We collected locality images for each specimen in this study using satellite data that was sourced from Bing VirtualEarth, and when this was unavailable from ESRI World Imagery (34, 35). In each case, we selected a square plot surrounding the collection event’s coordinates, 100 m on each side, at a scale of 1:2000. We visually inspected the images to exclude erroneous localities and crop out any human-made or modified structures. We reduced each image to its 8 most prevalent colors using kmeans clustering and extracted RGB color values from these in pavo (36). For comparison in the same color system, we transformed plumage reflectance spectra to RGB values using pavo. We took Euclidean distances between patches and background colors in RGB space. Summarizing these as the minimum, maximum, mean, or weighted mean showed similar results (Figure S3).

While we used an RGB color system for this analysis as a necessity based on the satellite data available, future launches of hyperspectral sensors will hopefully enable modeling of this process in the avian visual system.

We used phylogenetic relationships and time-calibrated branch lengths from a supermatrix phylogeny assembled by (37) using five mitochondrial and four nuclear markers. We pruned this tree to include only species for which color data was available (see Specimen Appendix for complete list).

We estimated the degree of evolutionary covariance in plumage coloration between different feather patches using a phylogenetically corrected Partial Least Squares (PLS) approach employed in a similar study (13). To do this, we used a two-block PLS regression method developed for use with highly dimensional traits and implemented in the R package geomorph (38). We performed hierarchical clustering (agglomerative nesting algorithm) with the package cluster (39) to infer groups of co-evolving color patches. We used the covariance ratio in geomorph to test whether variation among modules exceeds variation within them, in comparison to a simulated null (12).

To estimate the rate of evolutionary change in coloration, we used a method developed for highly dimensional traits (40). We applied this method to the first 10 PC axes of color to produce a joint rate estimate for each feather patch. To compare lightness differences between dorsal and ventral plumage patches, we used a phylogenetically corrected paired t-test in the R package phytools (41). For comparisons of NDVI and feather lightness, we used PGLS as implemented in the package caper (21), with likelihood-optimized lambda parameter. We compared distances between patches and satellite images across modules using phylogenetic generalized linear mixed models in the phyr package (22). Using this approach, we specified a model that included random effect terms for spatial and phylogenetic covariance matrices, as well as terms to remove potential pseudo-replication from species replicates and any duplicate localities.

## Supporting information

Specimen Appendix

## Acknowledgments

This study was inspired by conversations between NRF and CM Eliason at Evolution 2017. We thank Kaspar Delhey for his helpful comments on the manuscript. VR was supported by Palacky University, and NRF was supported by subsidy funding to OIST. We are eternally grateful to Leo Joseph, Robert Palmer, Margaret Causey and Alex Drew of the CSIRO Australian National Wildlife Collection, (ror.org/059mabc80) for their assistance in undertaking this research.

## Notes

### Competing Interest Statement

The authors have declared no competing interest.

