## Supplementary material for "Plumage color evolves as distinct dorsal and ventral modules in Australasian honeyeaters": Figure S1

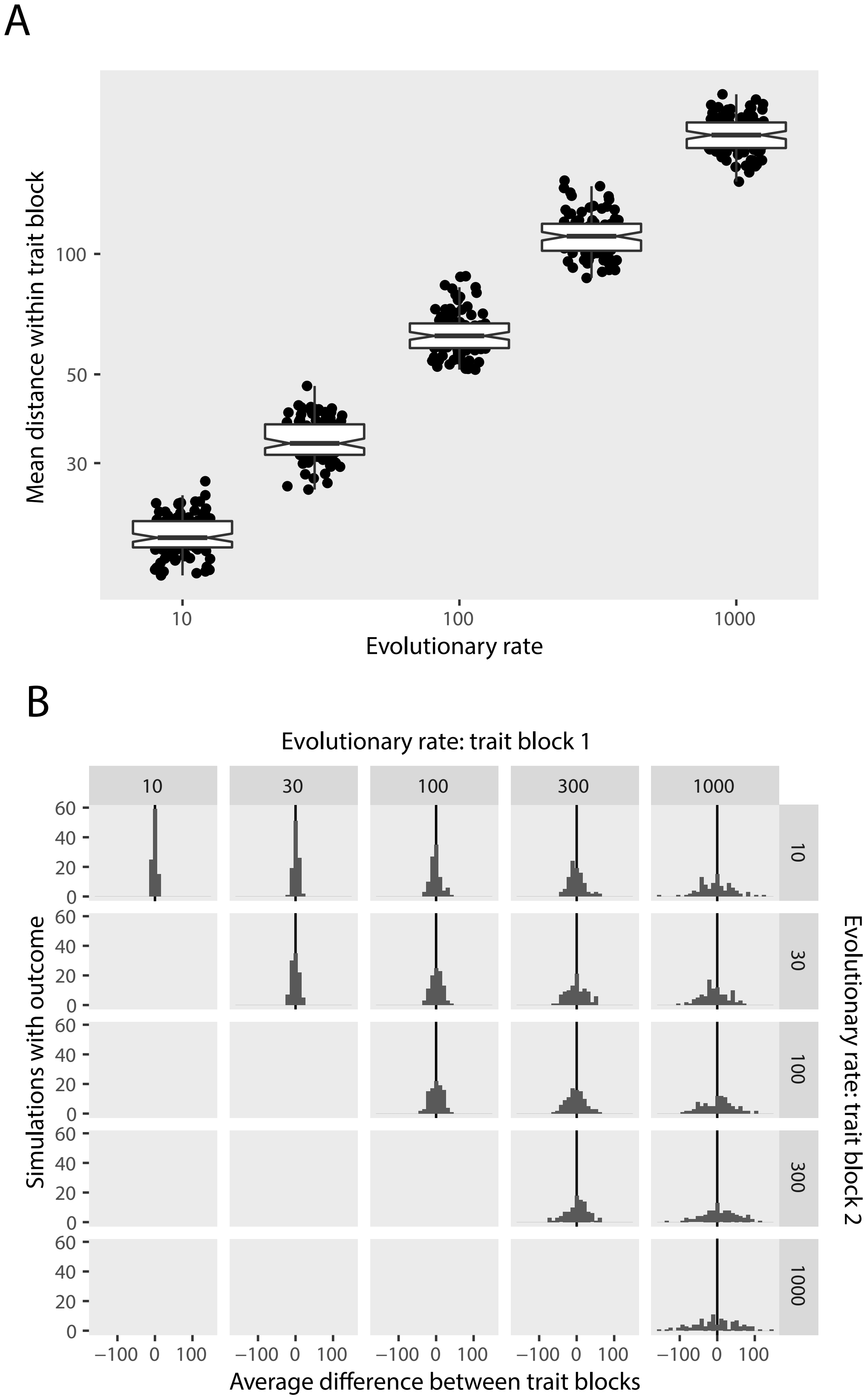


Figure S1: We used Brownian motion models implemented in the R package phytools to simulate blocks of 5 independently evolving traits on the Meliphagoidea phylogeny. Rates match the range described in this study. As the evolutionary rate of a trait block is increased, the internal contrast among traits within the block also increases (A). When multiple blocks are compared, rate differences are sufficient to produce contrast in many cases, however these are inconsistent in their direction. We interpret the implication of this result to be that high rates alone can explain internal contrast among ventral plumage patches, but cannot explain dorsoventral contrast where the dorsal side is consistenly darker than the ventral.
