## Supplementary material for "Plumage color evolves as distinct dorsal and ventral modules in Australasian honeyeaters": Figure S2

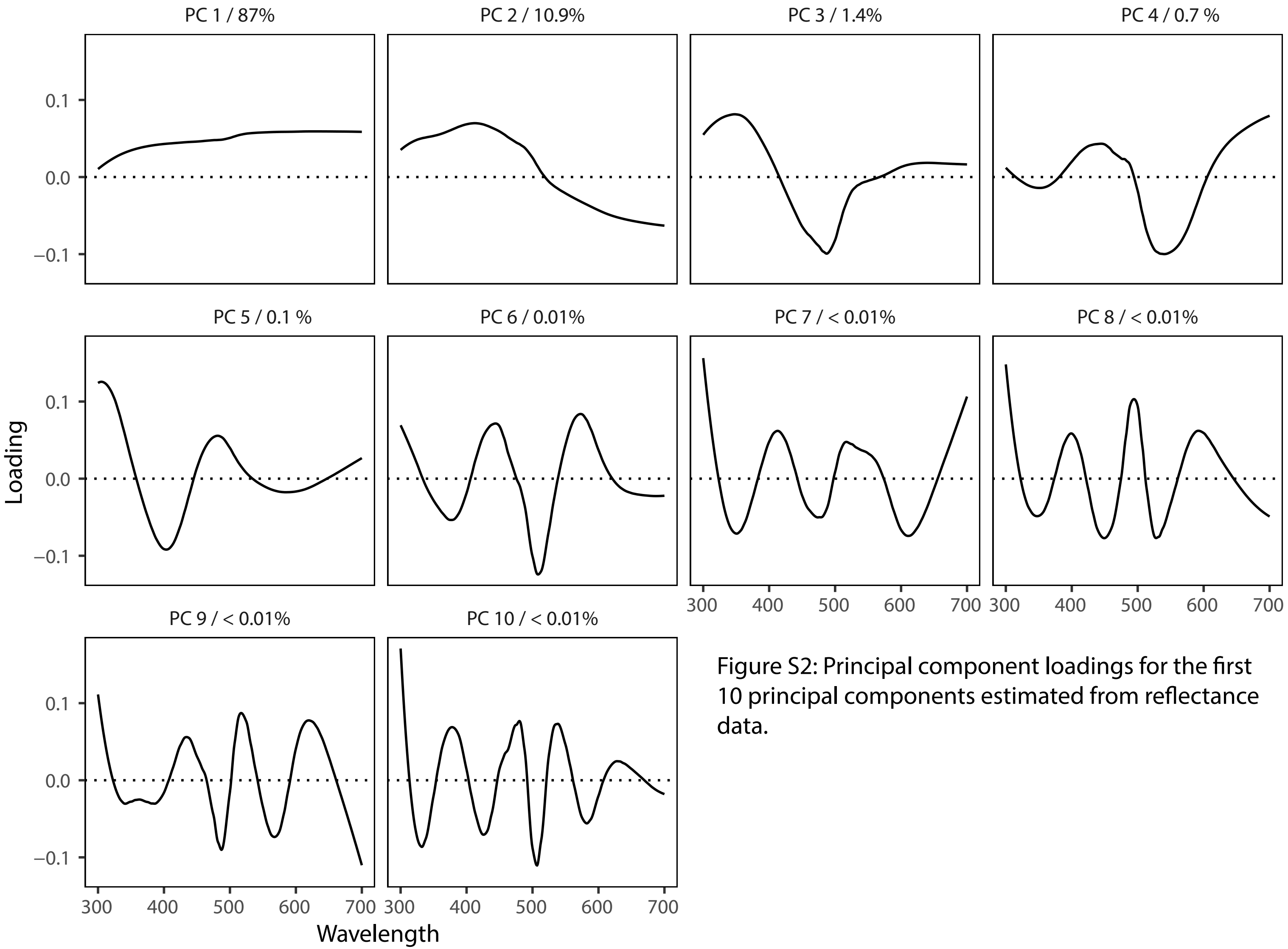


Figure S2: Principal component loadings for the first 10 principal components estimated from reflectance data.
