## Supplementary material for "Plumage color evolves as distinct dorsal and ventral modules in Australasian honeyeaters": Figure S3

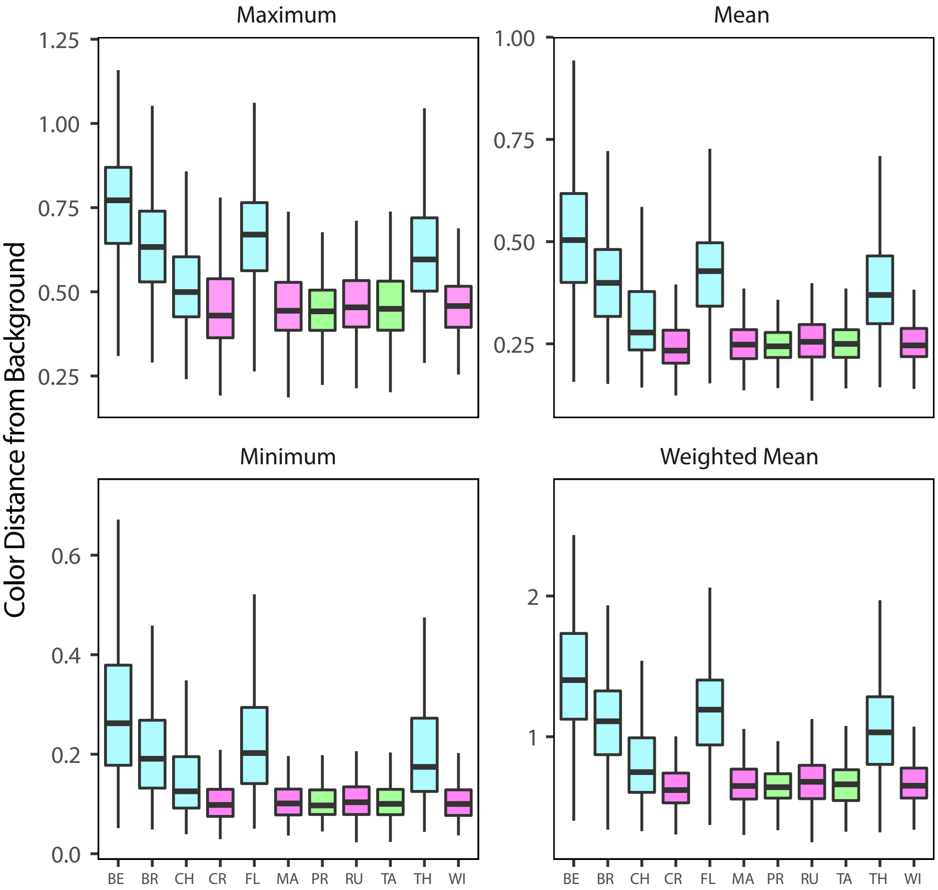


Figure S3: Color distances between feather patches and eight predominant colors from each specimen's sampling locality. These are summarized in four ways. The Maximum distance describes the distance between the feather color and the color of the most dissimilar of background colors. The minimum distance describes the distances between the feather color and the most similar background. The mean is the average distance between the feather color and all background colors, and the weighted mean includes a weighting factor that describes the proportion of pixels each background color occupies.
